# Structure of the enterococcal T4SS protein PrgL reveals unique dimerization interface in the VirB8 protein family

**DOI:** 10.1101/2020.10.30.342212

**Authors:** Franziska Jäger, Anaïs Lamy, Nina Guerini, Wei-Sheng Sun, Ronnie P-A Berntsson

**Affiliations:** Department of Medical Biochemistry and Biophysics, Umeå University, SE-90187 Umeå, Sweden; Wallenberg Centre for Molecular Medicine, Umeå University, Umeå, Sweden

**Author notes:** Correspondence should be addressed to R.P-A.B.

## Abstract

Multidrug resistant bacteria are one of the most important current threats to public health and a serious problem in hospital acquired infections (HAIs). Most antibiotic resistance genes are acquired via conjugative gene transfer, in a process that is mediated by a protein machinery called the Type 4 Secretion System (T4SS). The core of the T4SS is a multiprotein complex that spans both the cell wall and cellular membrane(s), serving as a channel for macromolecular secretion. Although the majority of multidrug resistant bacteria responsible for HAIs are of Gram-positive origin, with Enterococci being major contributors, mostly Gram-negative T4SSs have been characterized. Here we describe the structure and organisation of PrgL, one of the seven membrane proteins forming the translocation channel of the T4SS encoded by the pCF10 plasmid from *Enterococcus faecalis*. We present the structure of the C-terminal domain of PrgL, which displays similarity to VirB8 proteins of Gram-negative secretion systems. PrgL forms dimers and higher order oligomers but does not interact strongly with the other T4SS components. *In vitro* experiments show that the soluble domain alone is enough to drive both dimerization and dodecamerisation, with a dimerization interface that differs from all other known VirB8-like proteins. Our findings provide insight into the molecular building blocks of Gram-positive T4SS, highlighting similarities but also unique features in PrgL compared to other VirB8-like proteins.

## Introduction

Type 4 Secretion Systems (T4SS) are highly versatile molecular machines that are found in many bacterial species. This versatility is demonstrated by the fact that T4SSs take part in various functions such as i) transfer of DNA from bacterial donor cells into recipient cells, ii) transfer of effector proteins into eukaryotic cells, iii) exchange DNA with the milieu and iv) deliver of toxins to kill competing bacterial cells (Grohmann *et al.*, 2018). Over the past decade our understanding of the structure and function of T4SSs has increased drastically, with biochemical studies coupled to crystallography and cryo-electron microscopy (both single particle and tomography) of T4SS model systems (Peña *et al.*, 2012; Cascales *et al.*, 2013; Low *et al.*, 2014; Hu *et al.*, 2019; Hu, Khara and Christie, 2019). However, all well studied systems to date are of Gram-negative origin. Although the overall composition and structure of Gram-positive T4SSs deviate from their Gram-negative counterparts, we know rather little of them. For the conjugative T4SSs, the DNA processing proteins and the ATPases that drive transport are thought to be similar. However, the makeup of the actual T4SS channel is completely different, owing to Gram-positives only having one membrane and a thick cell-wall, compared to the Gram-negative double membrane and periplasmic space. Currently available knowledge of Gram-positive T4SSs are well summarized in previous reviews (Goessweiner-Mohr, Arends, *et al.*, 2013; Grohmann *et al.*, 2018).

Since its discovery at the end of the 1970’s, the conjugative plasmid pCF10 from *Enterococcus faecalis* has been serving as a model system for pheromone regulated conjugation in Gram-positive bacteria (Dunny, Brown and Clewell, 1978). The *pQ* promotor in pCF10 transcribes for one large operon that contains, among other things, all open reading frames for the entire T4SS (Dunny, 2013; Dunny and Berntsson, 2016). Previous research on components of this system have focussed on the DNA processing proteins and the extracellular adhesion proteins (Chen *et al.*, 2008; Li *et al.*, 2012; Bhatty *et al.*, 2015; Schmitt *et al.*, 2018, 2020; Rehman *et al.*, 2019). However, little information is currently available about the composition or architecture of the translocation channel. The current mechanistic model is that the accessory factor PcfF binds to the *oriT* of pCF10, where it then recruits the relaxase PcfG that nicks, unwinds and covalently binds to the ssDNA (Rehman *et al.*, 2019). This complex, called the relaxosome, is thought to recruit the T4 coupling protein PcfC, which initiates and provides energy for the DNA transfer through the translocation channel. (Chen *et al.*, 2008; Álvarez-Rodríguez *et al.*, 2020). The translocation channel itself consists of seven proteins (PrgD, -F, -H, -I, -K, -L and PcfH), each containing at least one transmembrane spanning helix (Alvarez-Martinez and Christie, 2009). These proteins are thought to form the membrane channel for export of the pCF10 transfer intermediate. However, no structural detail is available for any of these proteins, nor is the stoichiometry of these proteins in the channel known.

Two of the seven channel-forming proteins, PrgL and PrgD, have been predicted to have a VirB8-like fold (Goessweiner-Mohr, Grumet, *et al.*, 2013). In the Gram-negative systems, VirB8 proteins contain a single transmembrane helix and a larger soluble domain that goes by the name nuclear transport factor-2 (NTF2) domain, which is localized to the periplasmic space (Kumar and Das, 2001; Christie, 2016). VirB8 proteins are generally thought to form oligomers in the presence of their transmembrane domain, albeit examples exist of the NTF2 domain oligomerising on its own, such as the VirB8 homologue from the pKM101 system, TraE, which has been shown to form hexamers (Bourg *et al.*, 2009; Goessweiner-Mohr, Grumet, *et al.*, 2013; Fercher *et al.*, 2016; Casu *et al.*, 2018). VirB8 homologs have in numerous cases been shown to be essential for the function of their T4SS, and are suggested to play a role in complex assembly, but even in the well characterized Gram-negative systems, their structural roles have not yet been fully elucidated. Currently available subassemblies of Gram-negative T4SSs do not include VirB8, and thus provide no information on how VirB8 is part of the channel (Low *et al.*, 2014; Kwak *et al.*, 2017; Durie *et al.*, 2020; Sheedlo *et al.*, 2020).

In this study, we present the crystal structure of the soluble NTF2 domain of the VirB8 homologue PrgL from pCF10, and the biochemical and biophysical characterization of both its NTF2 domain and the full-length protein. Based on the presented data we propose that PrgL forms a cytoplasmic scaffold to facilitate the T4SS channel formation through the plasma membrane and peptidoglycan layer in *Enterococcus faecalis*.

## Results

Full-length PrgL is predicted to be a membrane protein with a short N-terminal domain followed by a single transmembrane (TM) helix and a large C-terminal domain. We utilized *Lactococcus lactis* as an expression host for overproduction of PrgL. *L. lactis* is closely related to *E. faecalis*, with both organisms belonging to the group of lactic acid bacteria (*Lactobacillales)*. Furthermore, *L. lactis* has been proven to be an effective host for membrane protein production in numerous cases (King, Boes and Kunji, 2015). We made a PrgL construct using GFP fused to the C-terminus as a folding indicator to measure protein expression via in-gel fluorescence.

SDS-PAGE in-gel fluorescence of whole cells after expression in *L. lactis* showed bands corresponding to the expected size of PrgL-GFP (**Fig 1a**). Since GFP only folds and becomes fluorescent in the reducing environment of the cytoplasm, this result indicates that PrgL has its C-terminus localised in the cytoplasm. (Feilmeier *et al.*, 2000). To further investigate this, we analysed the topology of N-terminally His-tagged PrgL (without GFP) in a protease protection assay using Proteinase K. The N-terminal His-tag of PrgL can be detected in the non-treated cell samples but not in the samples treated with proteinase K, either with or without cells lysis prior to the treatment **(Fig. 1b, right panel)**. In contrast, immunological detection of PrgL using an antibody raised against the soluble C-terminal domain of PrgL (PrgL_32-208_) is possible in the non-treated sample as well as in the Proteinase K treated sample and only disappears completely when the cells are lysed before the treatment **(Fig. 1b, left panel)**. Both the proteinase K assay and the GFP folding indicator points towards an N-out, C-in topology for PrgL, with the caveat that these findings are for PrgL on its own, expressed in *L. lactis*.

**Figure 1:**
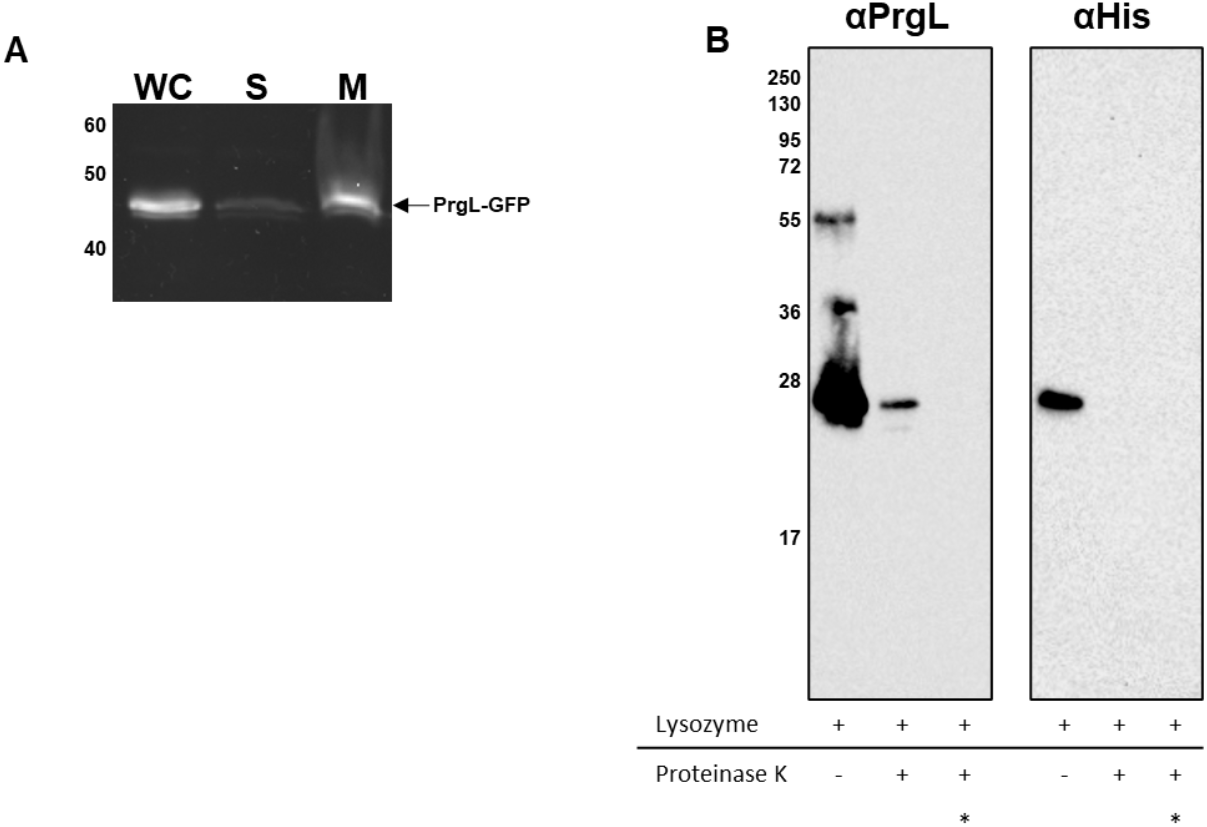
Cellular localization and topology of PrgL **A:** In-gel fluorescence of PrgL-GFP expressed in *L. lactis* NZ9000 cells harbouring recombinant PrgL. WC: whole cells; S: soluble fraction; M: membrane fraction. **B:** Analysis of the topology of PrgL. Samples were analysed by Western Blotting after treatment with proteinase K at a final concentration of 50 μg/ml. One sample was kept untreated as negative control. For the positive control, cells were lysed prior protease K treatment (lanes marked with an asterisk).

### PrgL forms both dimers and higher oligomers

Via in-gel fluorescence of PrgL-GFP, we then monitored the solubilization efficiency of nine different detergents (Bokman and Ward, 1981; Waldo *et al.*, 1999; Geertsma *et al.*, 2008). PrgL-GFP could be solubilized with all detergents apart from Lauryldimethylamineoxide (LDAO) (**Fig. 2a**). With in-gel fluorescence only monitoring correctly folded proteins, immunological detection of PrgL-GFP allowed us to determine the ratio between folded and misfolded protein (**Fig. 2b**). Extraction with octyl glucose neopentyl glycol (OGNG) gave the best result and was thus used for further purification of recombinant full-length PrgL-GFP via immobilised metal affinity chromatography (IMAC). Using size exclusion chromatography (SEC) we estimated the molar mass of PrgL-GFP to be approximately 160 kDa. (**Fig. S1a**). This apparent mass matches with the expected size of a full-length PrgL-GFP dimer (2 times 50.7 kDa) in the OGNG micelle, for which a molecular mass of 41 kDa has been reported (Casu *et al.*, 2018). Unfortunately the yield of full-length PrgL-GFP was too low to determine the exact molecular mass using size exclusion chromatography coupled to Multi-Angle-Light-Scattering (SEC-MALS) (Some *et al.*, 2019).

**Figure 2:**
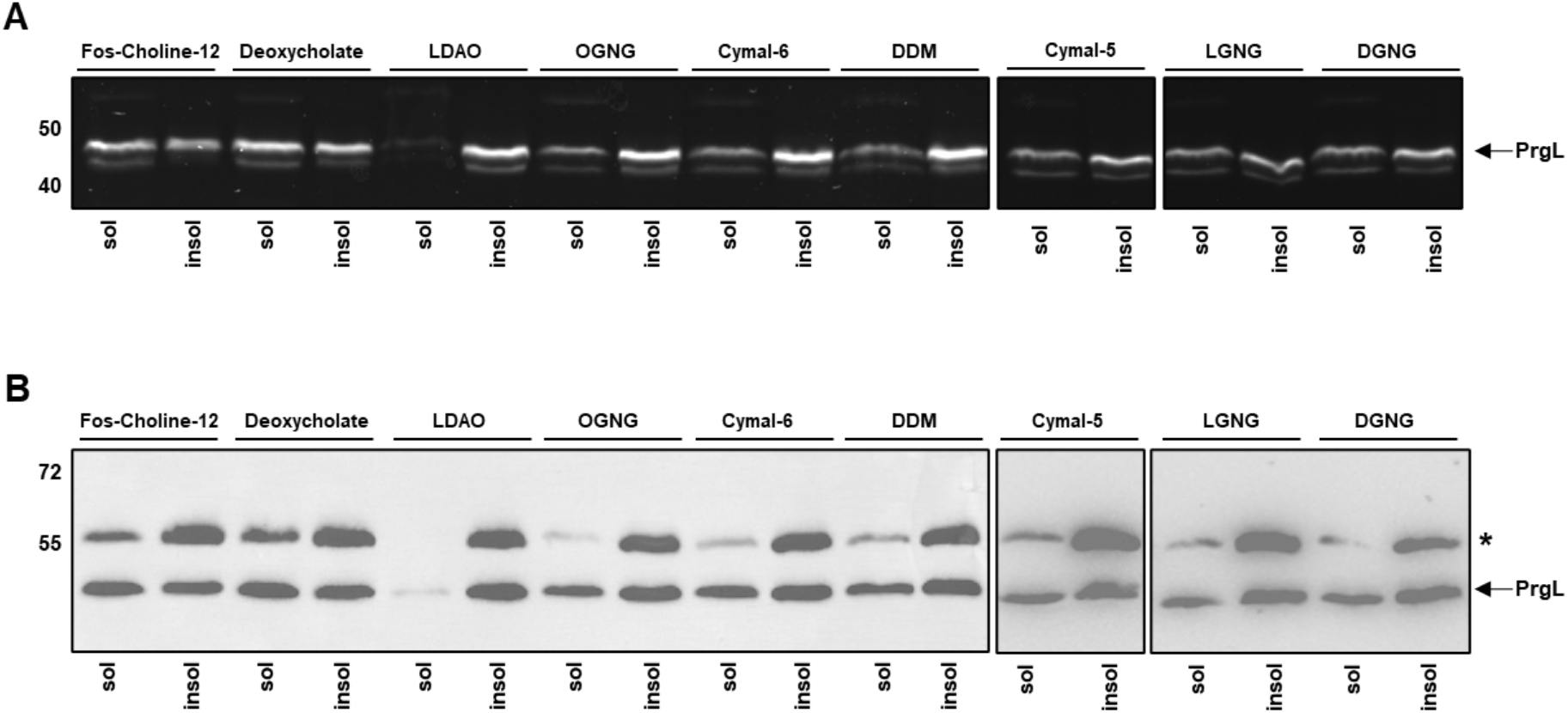
Detergent extraction of PrgL-GFP. Membrane fractions were treated with 1 % of the indicated detergent, as described under Experimental Procedures. **A:** About 10 μg of solubilised (sol) and insolubilised (insol) material were loaded on an SDS gel containing 12.5 % acrylamide. GFP fluorescence was visualised with help of the ChemiDoc™ Touch Imaging System from Bio-Rad using the CYBR Green setting. **B:** About 2 μg of solubilised (sol) and insolubilised (insol) material were loaded on an SDS gel containing 12.5 % acrylamide. Subsequently the proteins were transferred on a nitrocellulose membrane followed by immunological detection of the His-tag. Asterisk indicates misfolded PrgL.

As the yield of full-length PrgL proved difficult to scale up, we opted to determine the structure of the soluble domain of PrgL. PrgL_32-208_ was produced with a C-terminal 10x His-tag and purified with a yield of ca 50 mg/L culture. Purified PrgL_32-208_-His (M_W_ = 22.5 kDa) elutes as two separate peaks on SEC with the apparent masses of approximately 290 kDa and 55 kDa (**Fig. S1b**). SEC-MALS analysis shows that the later elution peak corresponded to a molecular mass of 43.7 +/− 0.4 kDa, which fits well with a dimeric form of PrgL_32-208_. The earlier peak was not as homogenous and showed a larger variation in the calculated MW between runs but had an average mass of 250 +/− 21.5 kDa, fitting with 10-12 copies of PrgL (**Fig. 3**). To verify whether this higher order oligomer is formed by full-length PrgL-His (M_W_ = 26 kDa) *in vivo*, we characterised the complex formation by *in vivo* crosslinking after recombinant expression in *L. lactis*. After treatment with paraformaldehyde (PFA) full-length PrgL-His migrated as a dimer and higher order oligomers on SDS-PAGE, with the largest complex migrating at the 250 kDa marker (**Fig. 4**). The 250 kDa elution peak of purified PrgL_32-208_-His was used for negative stain EM analysis where we observed ring-like particles of approximately 10 Å **(Fig. S2)**. Together with the crosslinking experiments the EM data and the SEC-MALS show that both PrgL_32-208_-His in solution and full-length PrgL-His *in vivo* form dimers and higher order oligomers. Our data show that the higher oligomer of PrgL contains up to 12 subunits, suggesting that it could be a dodecamer, similar to the stoichiometry which was observed of the VirB8 protein in the R388 T4SS from *E. coli* (Low *et al.*, 2014).

**Figure 3:**
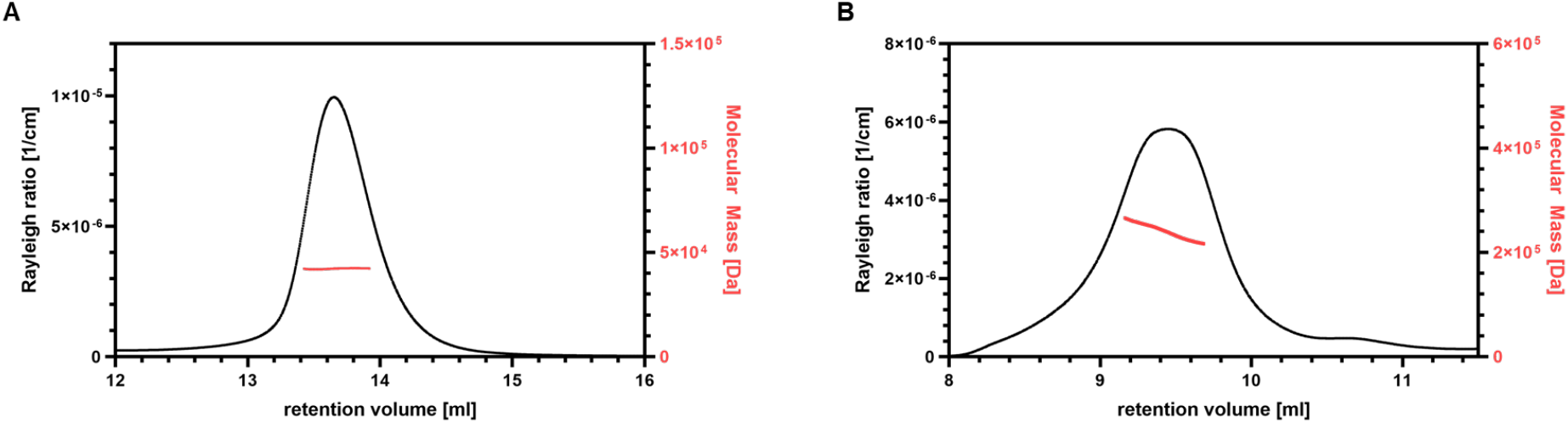
SEC-MALS of PrgL_32-208_. The elution profile of PrgL_32-208_ WT is shown with the average molecular mass of the dimer **(A)** and the higher order oligomer **(B)** calculated by MALS.

**Figure 4:**
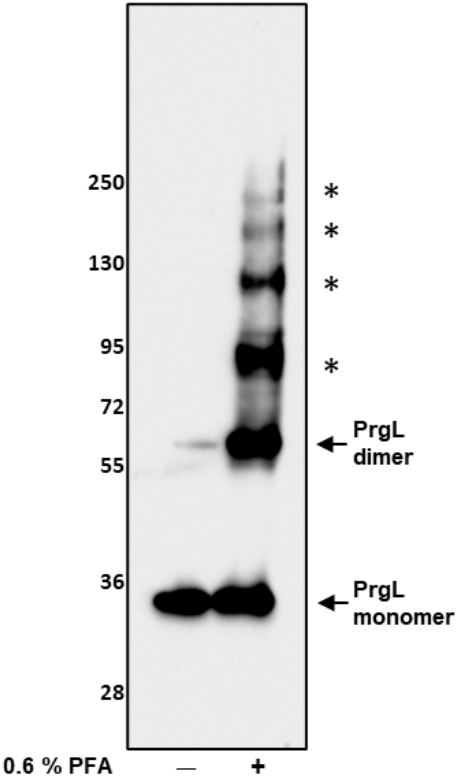
Analysis of the oligomerisation state of PrgL. Membrane fractions containing 5 μg of total amount of protein were analysed by Western Blotting after *in vivo* crosslinking of *L. lactis* whole cells with a final concentration of 0.6 % formaldehyde. Higher molecular mass complexes of PrgL formed after crosslinking are indicated with an asterisk (*).

### Structure of PrgL_32-208_ reveals its dimerization interface

Obtaining high quality cryo-EM grids with the PrgL_32-208_ dodecamer unfortunately proved challenging, as the protein complexes consistently fell apart upon vitrification. Instead we determined the structure of PrgL_32-208_ via X-ray crystallography using fractions from the dimeric SEC peak. The initial phases were derived from single-wavelength anomalous dispersion (SAD) experiments using selenomethionine-substituted protein and the final structure was determined to a resolution of 1.7 Å **(Table 1)**. The crystals belonged to the space group H3 and contained two molecules in the asymmetric unit. The final structural model corresponds to PrgL residues 52-189, as the N- and C-terminal ends were not visible in the electron density, likely due to them being flexible.

**Table 1:**
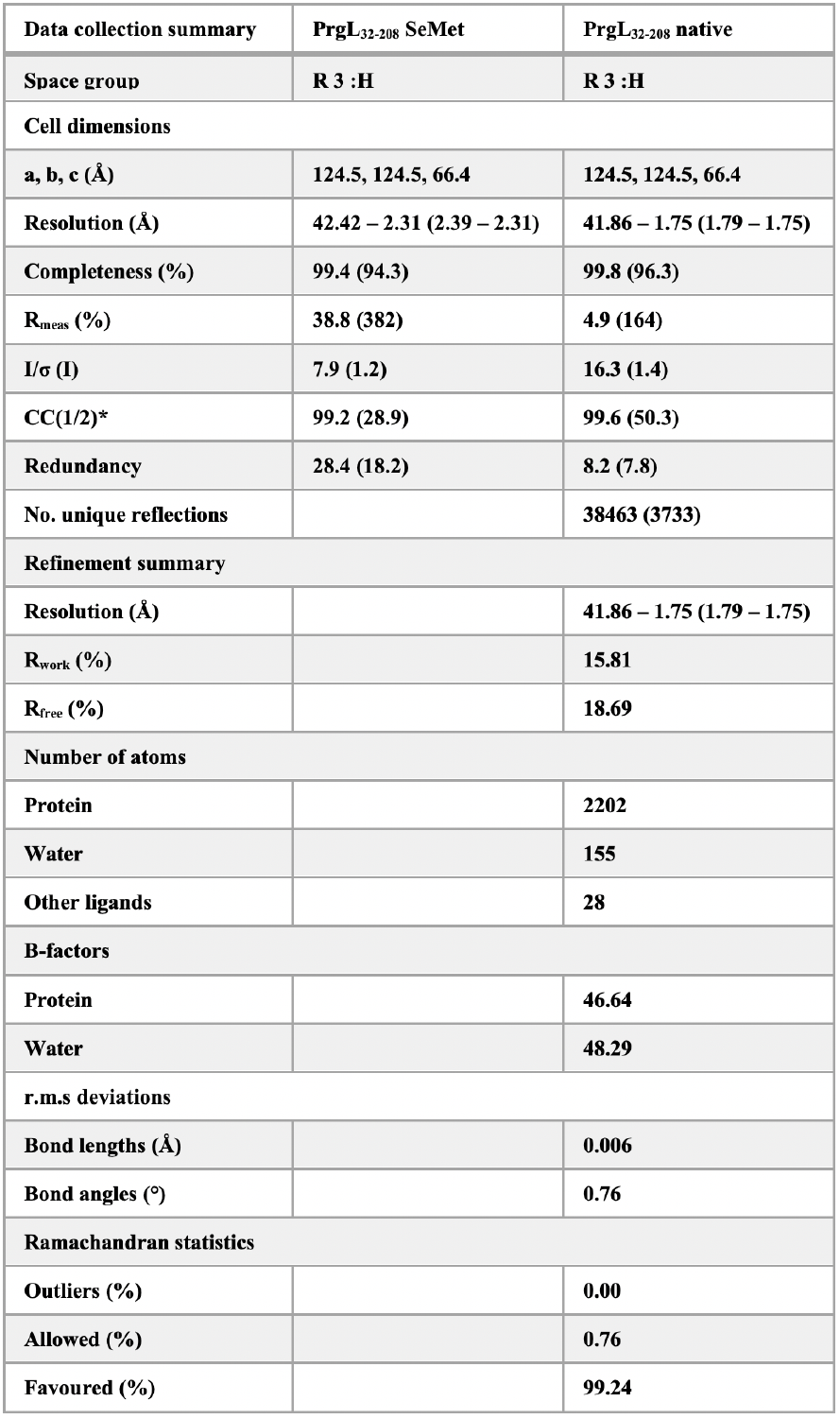
Data collection refinement statistics

| Data collection summary | PrgL<sub>32-208</sub> SeMet | PrgL<sub>32-208</sub> native |
| --- | --- | --- |
| Space group | R 3 :H | R 3 :H |
| Cell dimensions |  |  |
| a, b, c (Å) | 124.5, 124.5, 66.4 | 124.5, 124.5, 66.4 |
| Resolution (Å) | 42.42 – 2.31 (2.39 – 2.31) | 41.86 – 1.75 (1.79 – 1.75) |
| Completeness (%) | 99.4 (94.3) | 99.8 (96.3) |
| R<sub>meas</sub> (%) | 38.8 (382) | 4.9 (164) |
| I/σ (I) | 7.9 (1.2) | 16.3 (1.4) |
| CC(1/2)* | 99.2 (28.9) | 99.6 (50.3) |
| Redundancy | 28.4 (18.2) | 8.2 (7.8) |
| No. unique reflections |  | 38463 (3733) |
| Refinement summary |  |  |
| Resolution (Å) |  | 41.86 – 1.75 (1.79 – 1.75) |
| R<sub>work</sub> (%) |  | 15.81 |
| R<sub>free</sub> (%) |  | 18.69 |
| Number of atoms |  |  |
| Protein |  | 2202 |
| Water |  | 155 |
| Other ligands |  | 28 |
| B-factors |  |  |
| Protein |  | 46.64 |
| Water |  | 48.29 |
| r.m.s deviations |  |  |
| Bond lengths (Å) |  | 0.006 |
| Bond angles (°) |  | 0.76 |
| Ramachandran statistics |  |  |
| Outliers (%) |  | 0.00 |
| Allowed (%) |  | 0.76 |
| Favoured (%) |  | 99.24 |

The overall structure of PrgL_32-208_ consists of three antiparallel α-helices at the N-terminus and a highly curved β-sheet containing four antiparallel β-strands at the C-terminal end of the protein which is wrapped around the first helix (**Fig. 5a**). The third helix is kinked, and there is a twist in the fourth β-strand formed by Pro^167^. PrgL Tyr^66^ and Tyr^67^ form a negatively charged pocket in the centre of the protein, which has a Bis-Tris molecule (from the crystallisation conditions) bound. (**Fig. S3a +b**).

**Figure 5.**
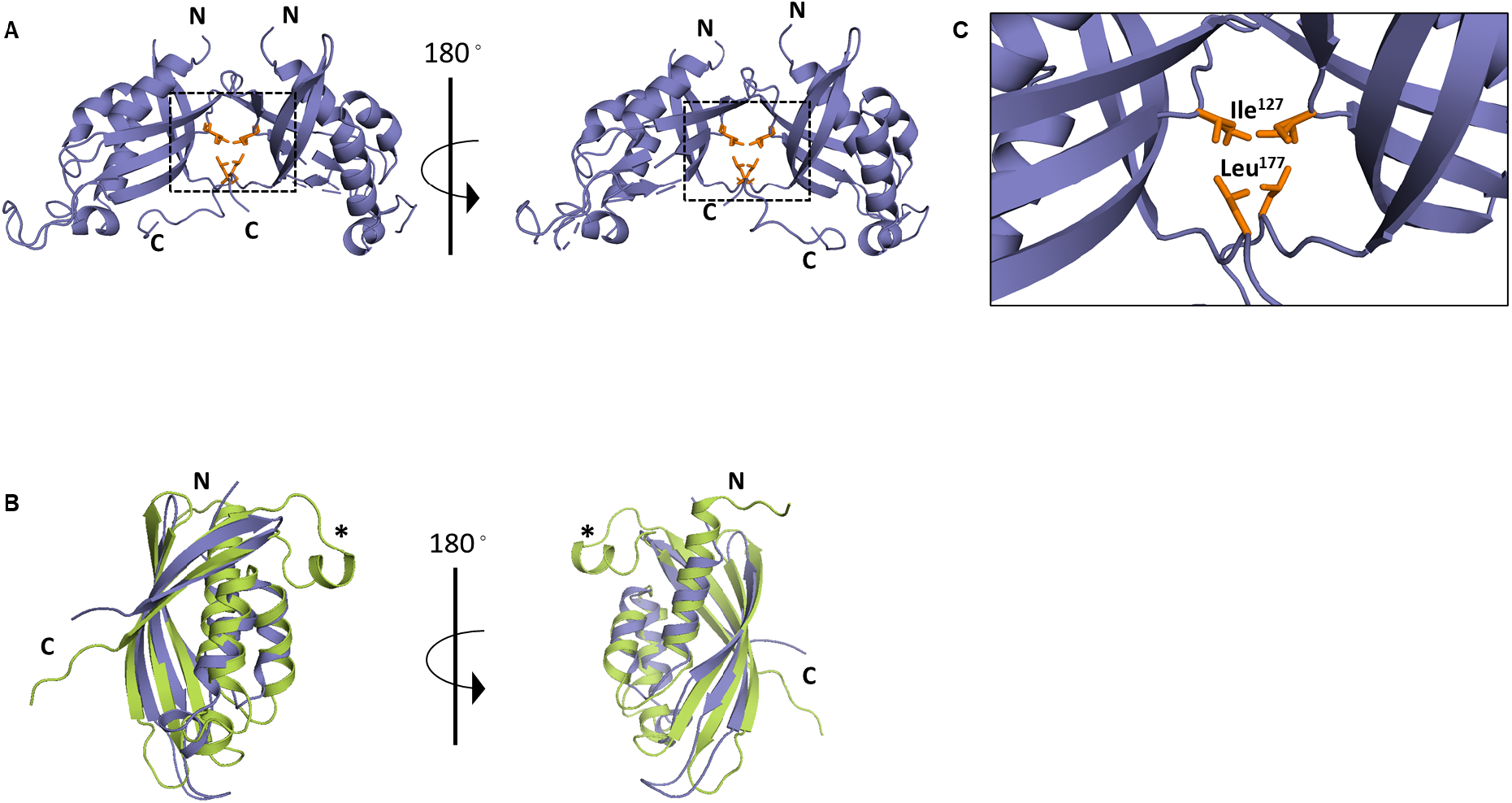
Crystal structure of PrgL_32-208_ at 1.7 Å resolution. **A:** Cartoon representation with PrgL coloured in blue, residues involved in the dimerization coloured in orange. **B:** Structural superimposition of PrgL_32-208_ (blue) and VirB8 from *A. tumefaciens* (PDB code: 2CC3, green). The extended loop with a short α-helix, found in nearly all VirB8-like proteins but absent in PrgL is marked with an asterisk (*). **C:** Enlarged view of the box from panels A, with the residues thought to be involved in dimerization highlighted in orange.

A DALI search (Holm, 2020) with the structure of PrgL_32-208_ confirms that PrgL is related to proteins from the VirB8 family, as it has the characteristic NTF2-like fold (**Table S1**). Compared to the archetypical VirB8 protein of *A. tumefaciens* (PDB code: 2CC3), PrgL is missing an extended loop and short α-helix between β3 and β4. This was also found to be the case for other VirB8-like proteins from *E. faecalis*: TraH (PDB code: 5AIW) and TraM (PDB code: 4EC6) of the pIP501 plasmid, and from other Gram-positive bacteria like *C. perfringens* (PDB code: 3UB1) **(Fig. 5b)**. PrgL_32-208_ can be superimposed with the two other VirB8 family proteins of *E. faecalis*: TraH and TraM with an r.m.s.d. value of 2.4 Å and 2.8 Å, respectively **(Table S1)**. Despite low sequence identity (12 – 14 %), the amino acids that form the negatively charged pocket in the centre of the PrgL (Tyr^66^ and Tyr^67^) are functionally conserved in both TraH (Phe^70^ and Tyr^71^) and TraM (Tyr^229^ and Phe^230^) **(Fig. S3c)**.

The soluble domain of various VirB8-like proteins is monomeric in solution. In contrast to this, PrgL_32-208_ forms both dimers and dodecamers in solution, as described above. Inspection of the PrgL_32-208_ structure revealed that the PrgL dimer interface consisted of a hydrophobic patch, with residues Ile^127^ and Leu^177^ (**Fig. 5c**). To test if this hydrophobic interface indeed mediated dimer formation, we mutated Ile^127^ to Glu. Purification of PrgL_32-208_:I127E shows a marked shift to a later elution volume on SEC (**Fig. 6a**), and SEC-MALS analysis showed that the peak fraction has a molecular mass of 26 ± 0.1 kDa, consistent with a monomeric state (**Fig. 6b**). The dimer interface seen in the crystal structure thus corresponds to the actual dimer interface in solution.

**Figure 6.**
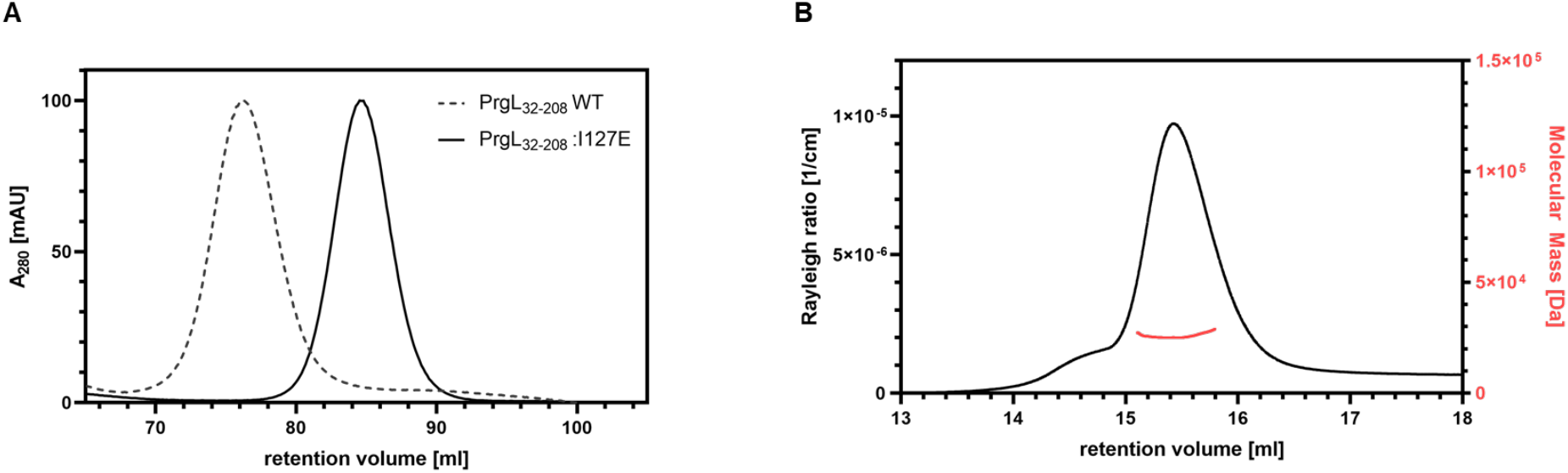
Elution profile of PrgL_32-208_ I127E. **A:** Elution profile of PrgL_32-208_:I127E in comparison to PrgL_32-208_ WT (dashed line) showing the later retention volume for the mutant. **B:** The elution profile of PrgL_32-208_:I127E is shown with the average molecular mass calculated by MALS.

### Interaction with other channel components

Since the Gram-positive T4SS channel assembly and the interaction between the different components remains unknown, we studied the interaction of PrgL with other components by *in vivo* crosslinking in *E. faecalis*. After inducing *E. faecalis* OG1RF:pCF10 cells with cCF10, cells were treated with PFA. Membrane fractions were prepared, loaded on SDS-PAGE and further analysed by Western Blot. Multiple bands could be detected with PrgL antibodies in the PFA cross-linked samples (**Fig. 7**). Comparing these higher mass bands with the ones obtained after crosslinking recombinantly expressed PrgL in *L. lactis*, the same pattern of bands could be detected, migrating at roughly the same molecular masses (**Suppl Fig. S4**). As the remaining T4SS machinery is absent in *L. lactis*, this indicates that PrgL does not strongly interact with other proteins of the T4SS or the *E. faecalis* membrane. Importantly, this experiment also validates the use of *L. lactis* as an expression host for PrgL, as it behaves and oligomerises in the same way in both *L. lactis* and *E. faecalis*.

**Figure 7.**
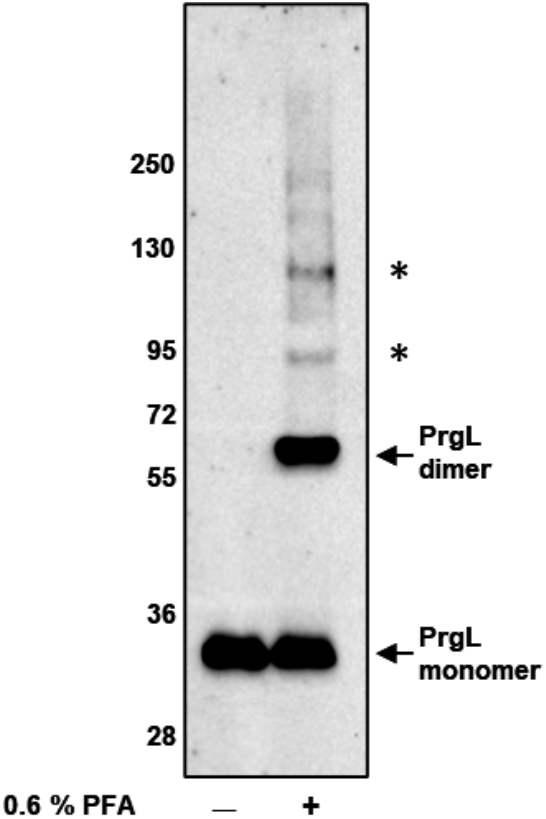
*In vivo* analysis of potential interaction of PrgL with other components of the conjugation channel. Membrane fractions containing 5 μg of total protein were analysed by Western Blotting after *in vivo* crosslinking of *E. faecalis* whole cells with a final concentration of 0.6 % formaldehyde.

## Discussion

Type 4 Secretion Systems are a main driver of horizontal gene transfer, including the spread of antibiotic resistance genes between bacterial species (Alvarez-Martinez and Christie, 2009). One of the best characterized model systems is the VirB/D4 T4SS from *A. tumefaciens,* which has been studied since the 1970s (Gurley *et al.*, 1979; Christie and Cascales, 2005; Li, Hu and Christie, 2019). Since then our understanding of these systems has deepened, and several systems have now structurally characterized the complete assembled T4SS (Bhatty, Laverde Gomez and Christie, 2013; Low *et al.*, 2014; Hu *et al.*, 2019). However, these characterized systems are all of Gram-negative origin. There is only limited biochemical or structural knowledge about T4SSs of Gram-positive origin, with the two systems that have been studied to some extent being the ones encoded on the pCF10 and pIP501 plasmids from *Enterococcus faecalis* (Kohler, Vaishampayan and Grohmann, 2018; Kohler, Keller and Grohmann, 2019; Sterling *et al.*, 2020).

In this study, we used structural, biophysical and biochemical approaches to characterise PrgL, a channel component of the T4SS from the conjugative plasmid pCF10 of *E. faecalis*. PrgL is predicted to be a membrane protein with a short N-terminal domain followed by a single transmembrane (TM) helix and a large C-terminal domain. Despite very low sequence identity with other T4SS transfer proteins of the VirB8 family **(Table S1)**, our results show that the overall structure of the C-terminal soluble domain of PrgL (PrgL_32-208_) has a similar nuclear transport factor-2 (NTF2)-like fold (Jakubowski, Krishnamoorthy and Christie, 2003) (**Fig. 5**).

VirB8-like proteins are known to oligomerise. A conserved NPXG motif, generally found on the extended loop between β3 and β4, is involved in the dimerization in nearly all VirB8 homologs described to date (Terradot *et al.*, 2005; Bailey *et al.*, 2006; Gillespie *et al.*, 2015). As PrgL is missing this extended loop, this motif is subsequently not present. The pIP501 protein TraH also lacks this motif, instead its oligomerisation dependents on its TM helix. We observed that PrgL oligomerization, into dimers and dodecamers, occurs in absence of either the NPXG motif or the TM helix (**Fig. 3**). Inspection of the dimeric PrgL_32-208_ structure revealed a hydrophobic pocket between the two monomers (**Fig. 5c**). A single amino acid substitution in this pocket, isoleucine to glutamic acid, resulted in only monomeric PrgL_32-208_ (**Fig. 6**). While it is well known that VirB8-like proteins like to crystallise as dimers or trimers, higher order oligomers were only described *in vivo* and for full length protein, making PrgL the first VirB8-like protein to multimerise *in vitro* by its NTF2-like domain alone, independent from the TM-helix or the NPXG motif (Terradot *et al.*, 2005; Bailey *et al.*, 2006; Fercher *et al.*, 2016).

According to the classification system for VirB8-like proteins, most Gram-positive variants belong to the BETA or GAMMA class (Goessweiner-Mohr, Grumet, *et al.*, 2013). Class BETA proteins contain two NTF2-like domains whereas GAMMA class VirB8-like proteins consist of a single C-terminal NTF2-like domain and a large cytoplasmic N-terminal domain. While class GAMMA-like proteins were exclusively found encoded on *E. faecalis* plasmids, PrgL is most closely related to classical Gram-negative VirB8 members belonging to the ALPHA class, which are characterised by harbouring only one NTF2-like domain. Prior our PrgL structure, only six proteins of Gram-positive origin were identified to belong to the ALPHA class, of which only the structure for TraH from the enterococcal pIP501 plasmid is available (Goessweiner-Mohr, Grumet, *et al.*, 2013; Fercher *et al.*, 2016). For VirB8-like proteins of all classes it has been shown that the NTF2-like domain resides outside the cell membrane, a characteristic that most Gram-negative and Gram-positive organisms seem to share (Terradot *et al.*, 2005; Bailey *et al.*, 2006; Porter *et al.*, 2012; Goessweiner-Mohr, Grumet, *et al.*, 2013). The only previously described VirB8 homolog that deviate from this rule is TraH from the pIP501 plasmid, which has been proposed to have its NTF2-like domain in the cytoplasm. Similar to TraH, our experiments using folding-indicator GFP and protease susceptibility indicate that PrgL also has its cytoplasmic NTF2-like domain in the cytoplasm (**Fig. 1**), at least when it is expressed on its own. In these experiments, lysozyme was used in order to hydrolyse the cell wall, as without lysozyme treatment no signal at all could be observed for either the anti-PrgL or anti-His antibodies. This treatment could make cells leaky, which might explain why we see a decrease of the PrgL C-terminal domain without previous cell lysis (**Fig. 1**). However, further validation is needed to confirm if PrgL has its NTF2 domain in the cytoplasm, and whether it is also the case when expressed in *E. faecalis* and in the presence of the rest of the T4SS components.

Although both PrgL and TraH_pIP501_ are from enterococcal plasmids, their T4SSs are quite dissimilar. pIP501 is a broad range conjugative plasmid, whereas pCF10 has a narrower range of recipients. The overall sequence similarity of their T4SSs is very low, and differences have also been reported with regards to their initial DNA processing (Kohler, Keller and Grohmann, 2019). TraH_pIP501is_ proposed to be a functional dimer, with the dimerization depending on the TM helix. PrgL on the other hand can form higher oligomers via its soluble domain, with dodecamers being observed for PrgL_32-208_ via SEC-MALS, as well as for the full-length PrgL *in vivo*. In this regard PrgL is so far unique, as other described soluble VirB8 domains do not form higher order oligomers in solution. That PrgL can form dodecamers after recombinant expression *in vivo*, in the absence of other T4SS components, could indicate that it is one of the early structural building blocks of the T4SS, similar to what has been proposed for TraH in its cognate pIP501 T4SS. Our *in vivo* cross-linking experiments on native pCF10 T4SS, expressed in *E. faecalis,* do not indicate strong interactions between PrgL and the other components. This resonates with Gram-negative VirB8 proteins, where the working hypothesis is that the interactions with other channel components are of a transient nature. VirB8-like proteins have been shown to be involved in the binding of the transferred DNA, either directly or via their binding partners, such as the relaxases (Cascales and Christie, 2004; Abajy *et al.*, 2007). Thus, it is possible that assembly of the T4SS machinery is depends upon DNA binding and processing, something that is not tested in our experimental set up.

The PrgL dodecamer has a ring-like structure, as visualized by our negative stain EM images (**Fig. S2**). We hypothesize that this ring-like structure can function as a scaffold to assemble other T4SS components into a functional channel. To our knowledge, this is the first time a Gram-positive VirB8 homologue has its higher order oligomer visualized. Also, for Gram-negative VirB8 homologs this high order oligomer has not been previously observed. The VirB8 from the pKM101 system has been shown to hexamerise. The R388 system has shown to contain 12 copies of VirB8, but whether these form an actual dodecamer, several smaller oligomers or are part of other heteromeric oligomers is not yet known. Whether this dodecamer formation is unique to PrgL or is more widespread remains to be determined.

To conclude, our data shows that PrgL has a VirB8 fold, validating previous bioinformatic studies (Bhatty, Laverde Gomez and Christie, 2013; Goessweiner-Mohr, Grumet, *et al.*, 2013). In contrast to its homologs, the soluble domain of PrgL is shown to form both dimers and dodecamers in solution, without aid of any NPXG motifs or its TM helix. The higher order oligomers are also observed with full-length PrgL *in situ*. Based on the available data of PrgL, and other VirB8 homologs, we propose that PrgL is involved in the early stages of the assembly process of the translocation channel by acting as a scaffolding protein for the other T4SS components. Experiments to determine the exact nature of PrgL in the assembly of the T4SS and its interaction with other channel components remain exciting avenues for future research.

## Material & Methods

### Bacterial strains and Plasmids

Plasmids encoding *prgL* variants for expression in *E. coli* and *L. lactis* were designed using the FXcloning system (Geertsma, 2014). Briefly, the *prgL* gene variants were PCR amplified from pCF10 using the complementary primers: 5’-ATATATGCTCTTCTAGTTATAATATGGAAAAATTAAGTAGATAC-3’ and 5’-TATATAGCTCTTCATGCCTTTTCCCCACTGTTTGCACTCTCCTT-3’ for full length PrgL or 5’-ATATATGCTCTTCTAGTAGTGTTGGGCAAAGAAAACAAGTAAAC-3’ and 5’-TATATAGCTCTTCATGCCTTTTCCCCACTGTTTGCACTCTCCTT-3’ for PrgL_32-208_, digested with SapI and cloned into the intermediate vector pINIT_kan. For expression in *L. lactis* the pINIT_PrgL variant was ligated into the pREXC3GH vector to generate a GFP and His-tagged variant while pINIT_PrgL_32-208_ was ligated into pREXC3H for a decahistidine-tagged variant of PrgL_32-208_ and the resulting plasmid transformed into *E. coli* TOP10 cells (Geertsma and Poolman, 2007). Subsequently, the *prgL* gene was transferred to pNZxLIC via SfiI digestion using the pSH71 harbouring vector pERL followed by transformation into *L. lactis* NZ9000. For expression in *E. coli* the PrgL_32-208_ variant was ligated into the p7XC3H before being transformed into *E. coli* BL21 cells.

For generating PrgL_32-208_ I127E, the pINIT_PrgL_32-208_ plasmid was PCR amplified from pINIT_PrgL_32-208_ using site-directed mutagenesis primers: 5’-GTC GAG GTG GAC GTC ACG TA-3’ and 5’-TC CAC CTC GAC TTC GTT GTT TTC-3’. Amplified products were digested with DpnI and subsequently transformed into *E. coli* strain TOP10. Further cloning for expression in *L. lactis* NZ9000 was followed after the same protocol as described above.

### Protein production and purification

PrgL variants were expressed as C-terminal decahistidine tagged proteins. For protein production in *L. lactis* NZ9000, cells were cultivated semianaerobically to an OD_600_ = 1.0 in 2 % Gistex (DSM Food Specialities B.V., Netherlands), 100 mM potassium phosphate (pH 7.0), 2.5 % (w/v) glucose, and 5 μg/ml chloramphenicol at 30 °C with gentle shaking before protein production was induced by adding 0.5 % (v/v) of culture supernatant of the nisin A producing strain *L. lactis* NZ9700 (de Ruyter, Kuipers and de Vos, 1996). Cells were pelleted 3 h after induction at 30 °C, flash-frozen in liquid nitrogen and stored at − 20 °C. For protein production of PrgL_32-208_ in *E. coli* BL21(DE3), cells were cultivated aerobically at 37 °C in TB medium supplemented with 0.4 % (v/v) Glycerol to an OD_600_ = 1.5, at which the temperature was lowered to 18 °C and production of the protein was induced by adding 0.4 mM IPTG. Cells were pelleted 16 h after induction, flash-frozen in liquid nitrogen and stored at −20 °C. Production of selenomethionine incorporated PrgL_32-208_ was carried out in *E. coli* BL21(DE3) grown in M9 minimal medium supplemented with 50 mg/L L-Selenomethionine as described earlier (Van Duyne *et al.*, 1993).

While full length PrgL resides in the membrane, PrgL_32-208_ is soluble and was purified from cytoplasmic extracts as follows: Cells were thawed and resuspended in Lysis Buffer 1 (20 mM HEPES/NaOH pH 7.8, 300 mM NaCl and 15 mM Imidazole pH 7.8). Cells were disrupted using a Constant cell disruptor (Constant Systems) by two or three passages at 25 000 psi or 39 000 psi when extracted from *E. coli* or *L. lactis*, respectively. Unbroken cells and cell debris were removed by centrifugation at 30 000 × *g* for 30 min. Proteins were purified at 4 °C on Ni-NTA-Sepharose (Macherey-Nagel) packed in a gravity flow column. The column was washed with a total of 30 column volumes (CV) of Wash Buffer (20 mM HEPES/NaOH pH 7.8, 300 mM NaCl, 50 mM Imidazole pH 7.8) including a 10 CV washing step with Wash Buffer supplemented with 2 M LiCl before bound proteins were eluted with Elution Buffer (20 mM HEPES/NaOH pH 7.8, 300 mM NaCl, and 500 mM Imidazole pH 7.8). Subsequently, the proteins were further purified using a Hiload 16/600 Superdex 200 pg gelfiltration column (GE Healthcare) in SEC Buffer (20 mM HEPES/NaOH pH7.8 and 250 mM NaCl). Two peaks eluting at the molecular mass corresponding to a dodecamer and dimer were observed on gel filtration. Both peak fractions were handled separately in the following steps. The protein was concentrated using Amicon Ultra Centrifugal Filters (Merk Millipore). Selenomethionine derivated PrgL_32-208_ was purified following the same protocol with the exception that 0.5 mM tris(2-carboxyethyl)-phosphine (TCEP) was added in all purification steps.

Full length PrgL was purified from membrane extracts as followed: Cells were thawed and resuspended in Lysis Buffer 2 (50 mM potassium phosphate pH 7.0, 10 % glycerol, and 1 mM MgSO_4_) before lysing them by three passages through a Constant cell disruptor (Constant Systems) at 39 000 psi. Unbroken cells and cell debris were separated by centrifugation at 30 000 × *g* for 30 min before membranes were collected by ultracentrifugation at 150 000 × *g* for 1 h. Subsequently, membranes were resuspended in Lysis Buffer 2 without MgSO_4_, flash-frozen in liquid nitrogen and stored at − 80 °C. Membrane-embedded proteins were solubilized by the addition of 1 % OGNG for 1.5 h at 4 °C in Solubilisation Buffer (50 mM potassium phosphate pH 7.8, 200 mM KCl, and 20 % glycerol). The solubilised proteins were separated from insoluble material by ultracentrifugation for 30 min at 150 000 × *g* and subsequently diluted by a factor of five in Solubilisation Buffer supplemented with 15 mM Imidazole pH 7.8) in order to reduce the detergent concentration. Proteins were purified as described above using Wash Buffer 2 (50 mM potassium phosphate pH 7.8, 200 mM KCl, 20 % glycerol, 0.2 % OGNG, and 50 mM imidazole pH 7.8 and eluted with Wash Buffer 2 supplemented with 500 mM Imidazole pH 7.8. Further purification of the proteins was achieved using a Superdex 200 increase 10/300 size exclusion column (GE Healthcare) in 50 mM HEPES/NaOH pH7.8, 150 mM NaCl, and 0.2 % OGNG.

### Size exclusion chromatography coupled to multi-angle light scattering (SEC-MALS)

Elution peaks corresponding to a molar mass of a dodecameric or dimeric protein of PrgL_32-208_ were further analysed by SEC-MALS with the use of an ÄKTApure system (GE Healthcare) coupled to a miniDAWN TREOS II detector and an OptiLab T-rEX online refractive index detector (Wyatt Technology). The absolute molar mass was calculated by analysing the scattering data using the ASTRA analysis software package, version 7.2.2.10 (Wyatt Technology). BSA was used for calibration and proteins were separated on a Superdex 200 Increase 10/300 analytical SEC column (GE Healthcare) with a flow rate of 0.4 ml/min. The dimer (4 mg/mL) and dodecamer peaks (2 mg/mL) were injected with 400 and 200 μL, respectively, and eluted in 20 mM HEPES/NaOH (pH 7.8) and 250 mM NaCl. The refractive index increment of PrgL_32-208_ was set at 0.185 ml/g and the extinction coefficient for UV detection at 280 nm was calculated from the primary structure of the protein construct.

### Electron Microscopy

For negative stain EM analyses 300 mesh carbon-coated copper grids were glow-discharged (PELCO easiGlow™) and incubated for 3 minutes with 3.5 μl SEC-purified sample at a final concentration of 10 ng/μl. Grids were subsequently stained with 1.5 % uranyl acetate and imaged at room temperature using an Talos L120C Transmission Electron Microscope (TEM).

### Crystallisation and structure determination

Native and selenomethionine derivatized PrgL_32-208_ were crystallised at 20 °C by sitting drop vapor diffusion in 0.1 M bis-Tris (pH 5.5), 25 % (w/v) PEG 3350 with a protein concentration of 6 mg/ml and a protein to reservoir ratio of 1:1. Crystals were flash-frozen in liquid nitrogen. X-ray diffraction data of native PrgL_32-208_ crystals were collected at beamline BioMAX at the MAX IV Laboratory (Lund, Sweden). Diffraction data of selenomethionine derivatised PrgL_32-208_ crystals were collected at beamline PX1 at the Swiss Light Source (SLS) (Paul Scherrer Institute, Switzerland). The data were processed using XDS (Kabsch *et al.*, 2010). The crystallographic phase problem was solved by means of single-wavelength anomalous dispersion (SAD), the selenomethionine sites were found and refined by Auto-Rickshaw pipeline and an initial model build by ARP/wARP (Panjikar *et al.*, 2005; Langer *et al.*, 2008). The crystals belong to space group H3 and contained two molecules in the asymmetric unit. Building of the model was performed in COOT and refined at 1.8 Å using PHENIX refine to R_work_ and R_free_ values of 17.9 % and 19.81 %, respectively (Adams *et al.*, 2002; Emsley and Cowtan, 2004). The structure has been deposited in the Protein Data Bank (PDB code: 7AED).

### In vivo crosslinking

For formaldehyde crosslinking in *L. lactis* NZ9000 full-length PrgL was produced in 1L of media as described above. 3h after induction 50 ml of the culture was harvested and cells resuspended in 5 ml 1 x PBS before being divided into two equal aliquots. Formaldehyde was added to one sample to a final concentration of 0.6 % while the second sample was kept untreated as negative control. Both samples were incubated at room temperature for 30 min before the reaction was quenched by addition of Tris/HCl (pH 8.0) to a final concentration of 100 mM. Cells were pelleted by centrifugation for 10 min at 3000 × *g,* resuspended in 20 ml Lysis buffer (50 mM potassium phosphate, pH 7.0, 10 % glycerol, 1 mM MgSO_4_) and treated as described above to acquire the isolated membrane fractions. Membranes were resuspended in 100 μl membrane buffer (50 mM potassium phosphate, pH 7.0, 10 % glycerol) prior to loading on SDS-PAGE and subsequent immunological detection of PrgL by Western Blotting with an anti-PrgL antiserum (1:10 000 dilution).

For interaction studies of PrgL with other proteins of the channel component the *E. faecalis* strain OG1RF harboring the pCF10 plasmid was cultivated in Brain heart infusion (BHI) media containing 10 μg/ml Tetracycline at 37 °C with gentle shaking to an OD_600_= 0.6-0.8 before induction with 5 ng/ml cCF10. Cells were pelleted after 1h statical incubation at 37 °C, washed in 50 ml 1 x PBS, pelleted again and subsequently resuspended in 6 ml crosslinking buffer (50 mM potassium phosphate, pH 7.0; 200 mM KCl; 250 mM Sucrose; 1 mM EDTA). One sample was incubated with PFA to a final concentration of 0.6%, while the other sample was kept untreated as negative control. Both samples were incubated at room temperature for 30 min before the reaction was quenched by addition of Tris/HCl (pH 8.0) to a final concentration of 100 mM. After a quenching time of 15 min samples were diluted with Lysis buffer (50 mM potassium phosphate, 10% glycerol, 1 mM MgSO_4_) to a total volume of 30 ml and treated as described for the purification of full-length PrgL to acquire the isolated membrane fractions. The membranes were resuspended in 200 μl of membrane buffer (50 mM potassium phosphate, pH 7.0, 10 % glycerol) prior to loading on SDS-PAGE and subsequent immunological detection of PrgL by Western Blotting with an anti-PrgL antiserum (1:10 000 dilution).

### Detergent solubilisation tests

For detergent solubilisation tests, membrane fractions from *L. lactis* were prepared as described above using a GFP-tagged variant of PrgL and 5 mg/ml of total protein were used for solubilisation. Samples were supplemented with the appropriate detergent to a final concentration of 1 % in 50 mM potassium phosphate (pH 7.0), 1 mM MgSO_4_, 10% Glycerol, and incubated for 1 h at 4 °C with gentle shaking. To separate the solubilised protein from the insoluble material, samples were centrifuged for 20 min at 89 000 × *g* at 4 °C. The supernatant was taken as the solubilised membrane protein and the pellet as the insoluble material. 10 μg of total protein was used for in-gel fluorescence and bands were visualised with help of the ChemiDoc™ Touch Imaging System from Bio-Rad. 2 μg of total protein was used for the immunological detection of PrgL by Western Blotting with an HRP-conjugated anti-His antiserum (1:10 000 dilution).

### Proteinase K digest

For proteinase K assays full-length N-terminal decahistidine tagged PrgL was produced in *L. lactis* NZ9000 as described above. After harvesting, cells were washed in 100 ml proteinase-buffer (30 mM Tris/HCl pH 7.5, 500 mM Sucrose, 10 mM CaCl_2_), pelleted for 20 min at 3200 × *g,* resuspended in 20 ml proteinase-buffer and divided into four equal aliquots à 5 ml before being flash-frozen and stored at −80 °C. For the proteinase K assay 5 ml of cells were thawed and aliquoted into 1 ml samples. Cells were treated with Lysozyme to pre-digest the cell wall at a final concentration of 30 mg/ml for 30 min at 30 °C. Following the Lysozyme treatment cells for the positive control samples were broken with the help of glass beads of 0.5 mm diameter (Sigma) using an Eppendorf shaker block at 4 °C and 18000 rpm for 20 min. Proteinase K digest was started by adding proteinase K to all samples with a final concentration of 50 μg/ml and incubating them for 30 min at 30 °C. The reaction was stopped by adding an EDTA free protease inhibitor tablet prior to the immunological detection of PrgL by Western Blotting with an anti-PrgL antiserum (1:10 000 dilution) and an anti-His antiserum (1:5 000 dilution).

## Acknowledgments

The authors thank Dr. Andreas Schmitt and Dr. Michael Järvå for help with the crystallographic data processing, Dr. Eric Geertsma for providing the plasmids of the FXCloning system, Dr. Josy ter Beek for critical reading of the manuscript and Prof. Peter J Christie and Prof. Gary Dunny for valuable discussions regarding the project. We are grateful to the beamline scientists at the MAX IV Laboratory (Lund, Sweden) for providing assistance in using beamline BioMax, as well as to the beamline scientist of beamline PX1 at the Swiss Light Source (Paul Scherrer Institute, Switzerland). This work was supported by grants from the Wenner-Gren Foundation (UPD2018-0008) to F.J, the Swedish Research Council (2016-03599), Knut and Alice Wallenberg Foundation, Kempestiftelserna (SMK-1762 & SMK-1869) and Carl-Tryggers stiftelse (CTS 18:39) to R.P-A.B.

## Author contributions (CRediT statement)

Franziska Jäger: Conceptualization, Investigation, Writing - Original Draft, Writing – Revision, Funding acquisition. Anaïs Lamy: Investigation, Validation. Nina Guerini: Investigation. Wei-Sheng Sun: Investigation. Ronnie Berntsson: Conceptualization, Writing - Original Draft, Writing – Revision, Supervision, Funding acquisition.

