## Supplementary materials for "Structure of the enterococcal T4SS protein PrgL reveals unique dimerization interface in the VirB8 protein family"

.

Franziska Jäger[1], Anaïs Lamy[1], Nina Guerini[1], Wei-Sheng Sun[1] and Ronnie P-A Berntsson[1,2]

[1] Department of Medical Biochemistry and Biophysics, Umeå University, SE-90187 Umeå, Sweden

[2] Wallenberg Centre for Molecular Medicine, Umeå University, Umeå, Sweden

**Table S1**: Structural alignment with DALI. The *E. faecalis* homologue proteins TraH and TraM are highlighted in blue.

| PDB | Z-score | RMSD [Å] | aligned residues | total residues | sequence identity [%] | Description | source | Gram [+/-] |
|---|---|---|---|---|---|---|---|---|
| 4jf8 | 12.9 | 3 | 119 | 144 | 12 | TrwG component of type IV secretion system | Bartonella birtlesii | negative |
| 4nhf | 12.8 | 2.8 | 119 | 142 | 7 | TrwG protein | Bartonella grahamii | negative |
| 5i97 | 12.6 | 3.2 | 119 | 137 | 8 | Conjugal transfer protein | Escherichia coli | negative |
| 5jbs | 12.5 | 3 | 116 | 136 | 9 | Type IV secretion system protein virB8 | Brucella suis | negative |
| 4o3v | 12.3 | 3.1 | 118 | 138 | 13 | VirB8-like protein of type IV secretion system | Rickettsia typhi | negative |
| 4lso | 12 | 2.7 | 117 | 141 | 12 | Type IV secretion system protein virB8 | Bartonella quintana | negative |
| 4mei | 12 | 2.7 | 116 | 136 | 9 | VirB8 protein | Bartonella tribocorum | negative |
| 5aiw | 11.9 | 2.4 | 112 | 127 | 14 | TraH | Enterococcus faecalis | positive |
| 4kz1 | 11.8 | 2.8 | 116 | 135 | 11 | VirB8 protein | Bartonella grahamii | negative |
| 2cc3 | 11.7 | 3 | 118 | 144 | 6 | VirB8 protein | Agrobacterium tumefaciens | negative |
| 5cnl | 11.6 | 2.6 | 116 | 139 | 10 | IcmL-like | Legionella pneumophila | negative |
| 3wz4 | 11.4 | 2.4 | 118 | 140 | 8 | DotI | Legionella pneumophila | negative |
| 6iqt | 11.3 | 3.1 | 118 | 135 | 10 | Cag pathogenicity island protein (Cag10) | Helicobacter pylori | negative |
| 5d9r | 10.9 | 3.6 | 116 | 132 | 7 | Protein accumulation and replication of chloroplasts 6, chloroplastic | Arabidopsis thaliana | plant |
| 3ub1 | 10.7 | 3 | 110 | 248 | 15 | ORF13-like protein | Clostridium perfringens | positive |
| 3wz3 | 10.7 | 2.8 | 116 | 137 | 8 | TraM protein | Plasmid R64 | negative |
| 4ec6 | 9.9 | 2.8 | 98 | 109 | 12 | TraM | Enterococcus faecalis | positive |

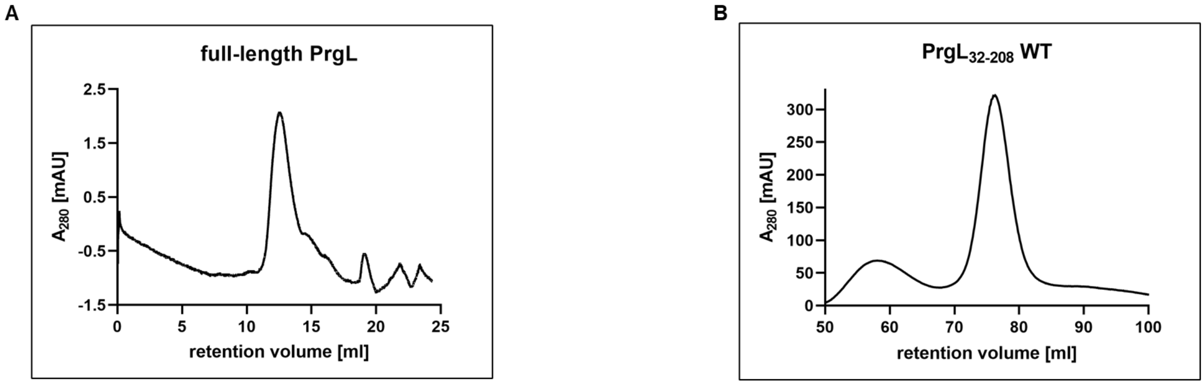


**Figure S 1**: SEC elution profile, monitored at 280 nm, of full-length PrgL (left panel, Superdex 200 Increase 10/300 GL column) and $PrgL_{32\text{-}208}$. (right panel, Superdex HiLoad 16/600 column)

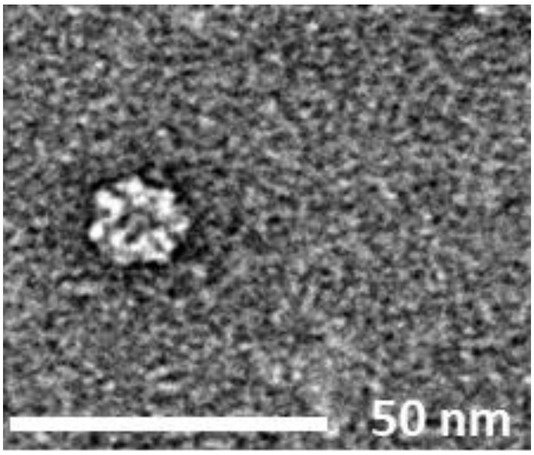


**Figure S2**: Electron microscopy analysis of $PrgL_{32\text{-}208}$. Negative-stain micrograph of the $PrgL_{32\text{-}208}$ complex showing a particle of ~10 Å.

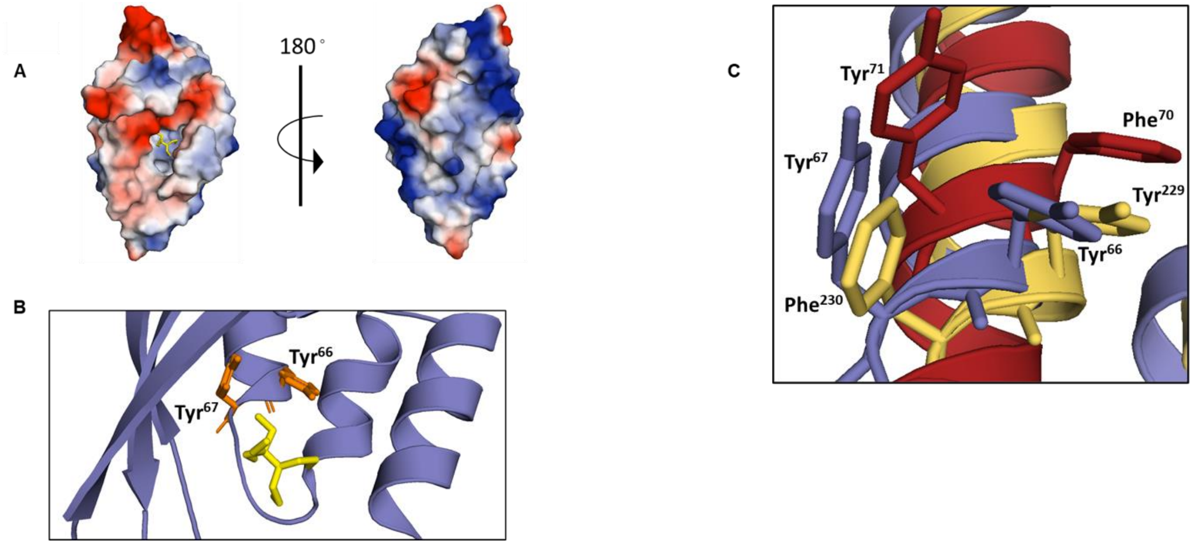


**Figure S3**: Crystal structure of the soluble domain PrgL$_{32-208}$ at 1.7 Å resolution. **A:** Surface charge distribution indicating a positive (red) and negative (blue) surface charge. A bound bis-Tris molecule within the negatively charged cleft is shown as sticks. **B:** Aromatic residues located in the negatively charged cleft facing the bound bis-Tris molecule. **C:** Enlarged view on the aromatic residues (purple) in the negatively charged cleft aligned with the structures of the *E. faecalis* homologues TraH (PDB code: 5AIW, red) and TraM (PDB code: 4EC6, yellow).

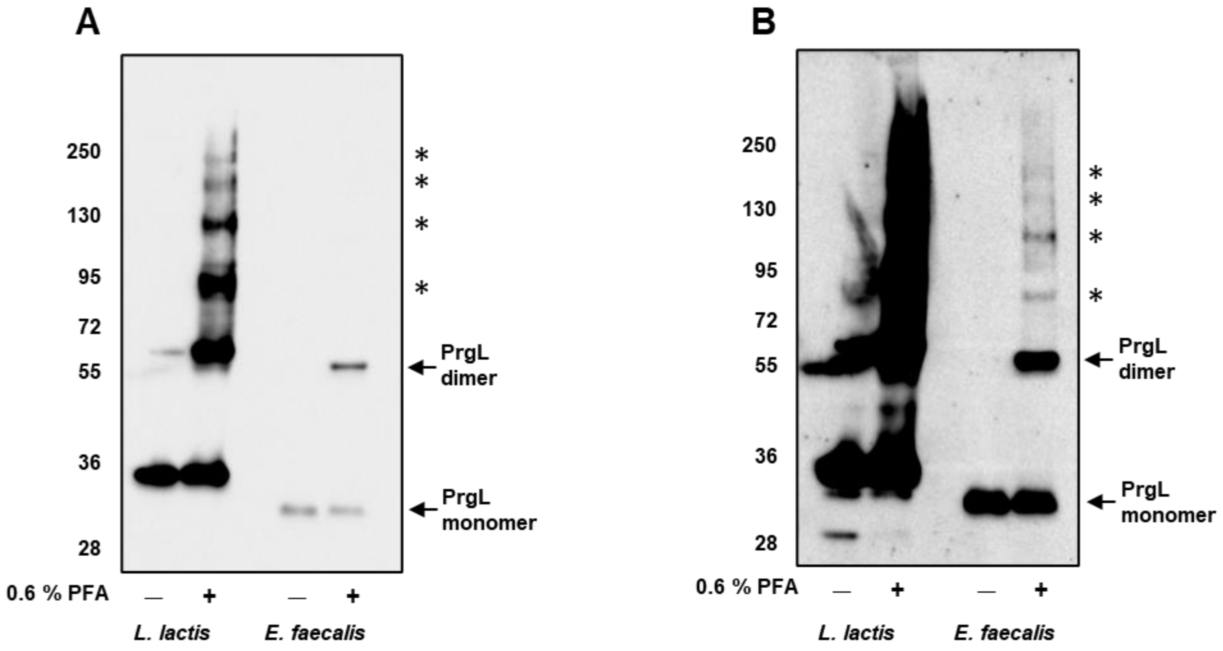


**Figure S4.** Migration pattern of full-length PrgL on a 4-20 % gradient gel. Shown is the Western Blot used for Fig. 4 and Fig. 7 with an exposure time of 5 sec (A – used for Fig. 4) and 5 min (B – used for Fig. 7). When expressed in *L. lactis*, PrgL is overexpressed as a recombinant variant carrying a decahistidine-tag and a 3C cleavage site. This means that the *L. lactis* PrgL monomer is 2 kDa larger than the native PrgL expressed from pCF10 in *E. faecalis*, explaining the differences in migration between the *L. lactis* and *E. faecalis* versions on the blots. Higher molecular mass complexes of PrgL formed after crosslinking are indicated with an asterisk (*).
